# Variance Adaptive Shrinkage (vash): Flexible Empirical Bayes estimation of variances

**DOI:** 10.1101/048660

**Authors:** Mengyin Lu, Matthew Stephens

**Affiliations:** Department of Statistics, University of Chicago; Department of Human Genetics, University of Chicago

## Abstract

**Motivation:** We consider the problem of estimating variances on a large number of “similar” units, when there are relatively few observations on each unit. This problem is important in genomics, for example, where it is often desired to estimate variances for thousands of genes (or some other genomic unit) from just a few measurements on each. A common approach to this problem is to use an Empirical Bayes (EB) method that assumes the variances among genes follow an inverse-gamma distribution. Here we describe a more flexible EB method, whose main assumption is that the distribution of the variances (or, as an alternative, the precisions) is unimodal.

**Results:** We show that this more flexible assumption provides competitive performance with existing methods when the variances truly come from an inverse-gamma distribution, and can outperform them when the distribution of the variances is more complex. In analyses of several human gene expression datasets from the Genotype Tissues Expression (GTEx) consortium, we find that our more flexible model often fits the data appreciably better than the single inverse gamma distribution. At the same time we find that, for variance estimation, the differences between methods is often small, suggesting that the simpler methods will often suffice in practice.

**Availability:** Our methods are implemented in an R package **vashr** available from http://github.com/mengyin/vashr.

## 1 Introduction

In statistics, shrinkage procedures are important ways to help improve estimation accuracy. They are often used to shrink target parameters towards some prior belief, and typically reduce the variance of the resulting estimators by borrowing information from prior knowledge or other observations. In this paper, we propose a flexible Bayesian shrinkage estimator for variances, which is able to reduce variance estimation error rates in some particular genomic settings.

Shrinkage estimators have been widely used in genomics, particularly, for example, to increase power to identify differentially expressed genes in two or more conditions. A typical pipeline for identifying differentially expressed genes computes a p-value for each gene using a t-test (two condition experiments) or F-test (multiple condition experiments), both of which require an estimate of the variance in expression of each gene among samples. In the classical t-test or F-test, sample variances are used as plugin estimates of gene-specific variances. However, when the sample size is small, sample variances can be inaccurate, resulting in loss of power [Murie et al., 2009]. Hence, many methods have been proposed to improve variance estimation. For example, several papers [Tusher *et al*., 2001, Efron *et al*., 2001, Broberg *et al*., 2003] suggested adding an offset standard deviation to stabilize small variance estimates. A more sophisticated approach [Baldi and Long, 2001] uses a parametric hierarchical model to combine information across genes, using an inverse gamma prior for the variances, and a Gamma likelihood to model the observed sample variances. This idea was further developed by Smyth [2004] into an Empirical Bayes approach that estimates the parameters of the prior distribution from the data. This improves performance by making the method more adaptive to the data. Smyth [2004] also introduces the “moderated t-test”, which modifies the classical t-test by replacing the gene-specific sample variances with estimates based on their posterior distribution. This pipeline, implemented in the software limma, is widely used in genomics thanks to its adaptivity, computational efficiency and ease of use.

While assuming an inverse-gamma distribution for the variances yields simple procedures, the actual distribution of variances may be more complex. Motivated by this, Phipson et *al.* [2013] (limma with robust option, denoted by ‘limmaR’) modified the procedures from Smyth [2004], in a somewhat *ad hoc* way, to allow that some small proportion of outlier genes may have higher variability than expected under the inverse-gamma assumption. They showed that, in the presence of such outliers, this procedure could improve on the standard limma pipeline.

Here we consider a more general and adaptive approach to this problem. Our method is based on the assumption that the distribution of the variances (or, alternatively, the precisions) is unimodal. We use a mixture of (possibly a large number of) inverse-gamma distributions to flexibly model this unimodal distribution, and provide simple computational procedures to fit this Empirical Bayes model by maximum likelihood of the mixture proportions. Our procedure provides a posterior distribution on each variance or precision, as well as point estimates (posterior mean). The methods are an analogue of the “adaptive shrinkage” methods for mean parameters introduced in Stephens [2016], and are implemented in the R package vashr (for “variance adaptive shrinkage in R”). We compare our method with both limma and limmaR in various simulation studies, and also assess its utility on real gene expression data.

## 2 Methods

### 2.1 Models

Suppose that we observe variance estimates 
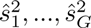
 that are estimates of underlying “true” variances 
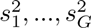
 Motivated by standard normal theory, we assume that

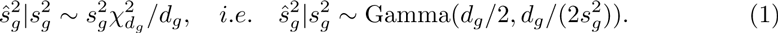

where the degrees of freedom *d_g_* depends on the sample size and we assume it to be known.

Empirical Bayes (EB) approaches to estimating 
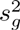
 (eg [Smyth, 2004]) are commonly used to improve accuracy, particularly when the degrees of freedom *d_g_* for each observation are modest. The EB approach typically assumes that the variances 
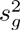
 are independent and identically distributed from some underlying parametric distribution *g*:

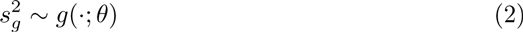

where the parameters *θ* are to be estimated from the data. Equivalently, that the precisions (inverse variances), 
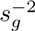
, are i.i.d. from some *h*(·; *θ*). A standard approach (Smyth) assumes that g is an inverse-gamma distribution (i.e. h is a gamma distribution) which simplifies inference because of conjugacy. Here we introduce more flexible assumptions for *g* or *h:* specifically that either *g* or *h* is *unimodal.* By using a mixture of inverse gamma distributions for *g* (i.e. a mixture of gamma distributions for *h*), we can flexibly capture a wide variety of unimodal distributions for *g* or *h*, while preserving many of the computational benefits of conjugacy.

### 2.2 A unimodal distribution for the variances

Let InvGamma(·;*a, b*) denote the density of an inverse-gamma distribution with shape *a* and rate *b*. This distribution is unimodal with mode at *c* = *b*/(*a* + 1). To obtain a more flexible family of unimodal distributions with mode at *c* we consider a mixture of inverse-gamma distributions, each with mode at *c*:

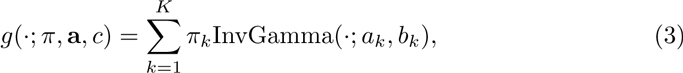

where

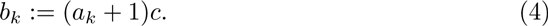

Each component in (3) has mode at *c*, and the variance about this mode is controlled by *a_k_*, with large *a_k_* corresponding to small variance. By setting **a** to a fixed grid of values that range from “small” to “large”, we obtain a flexible family of distributions, with hyperparameters *π*, that are unimodal about *c*. See below for details on choice of grid for **a**. This approach is analogous to Stephens [2016], which uses mixtures of normal or uniform distributions, with a fixed grid of variances, to model unimodal distributions for mean parameters.

### 2.3 Estimating hyper-parameters

For *K* = 1 we estimate the hyperparameters (*a*, *c)* by maximising the likelihood

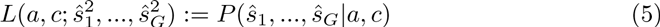

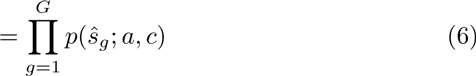

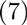

where

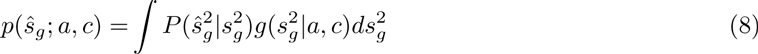

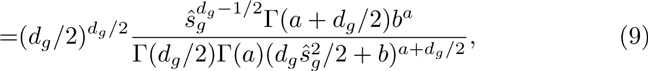

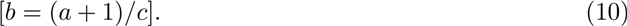

We use the R command optim to numerically maximize this likelihood. The approach is similar to Smyth [2004], except that we use maximum likelihood instead of moment matching.

For *K* > 1, as noted above, we use *K* “large” (e.g. 10-16), fix the values of *a_k_* to a grid of values from “small” to “large”, and estimate the hyper-parameters *c*, *π* by maximizing the likelihood

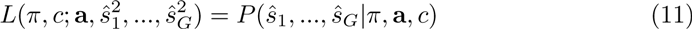

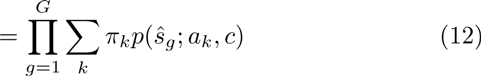

where *p*(*ŝ_g_*;*a_k_*,*c*) is given by (9). We center the grid of *a_k_* values on the point estimate *â* obtained for *K* = 1, to ensure that the grid values span a range consistent with the data. Moreover, if the data are consistent with *K* = 1 then the estimated *π* will be concentrated on the component with *a_k_* = *â*, and thus lead to similar results to limma.

To maximize the likelihood we use an iterative procedure that alternates between updating *c* and *π*, with each step increasing the likelihood. Given *c*, we update *π* using a simple EM step [Dempster *et al.*, 1977]. Given *π* we update *c* by optimizing (12) numerically using optim. We use SQUAREM [Varadhan, 2010] to accelerate convergence of the overall procedure. See Appendix for details.

### 2.4 Posterior calculations

Using (3) as a prior distribution for 
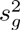
, and combining with the likelihood (1) the posterior distribution of 
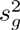
 is also a mixture of inverse-gamma distributions:

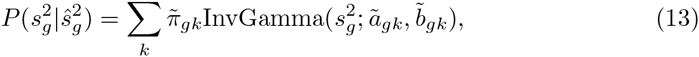

where

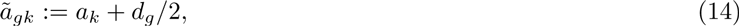

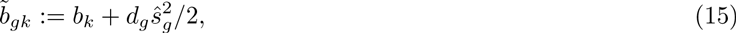

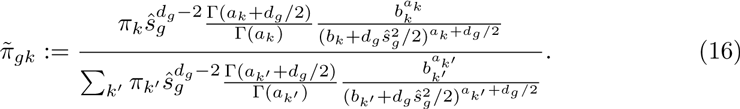

Following Smyth [2004] we use the posterior mean of 
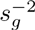
 as a point estimate for the precision 
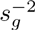
:

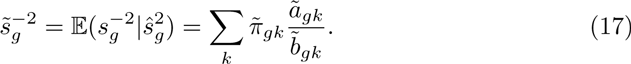

Note that each 
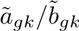
 can be interpreted as a shrinkage-based estimate of 
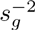
, since it lies between the observation 
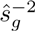
 and the prior mean of the *k*th mixture component *a_k_*/*b_k_*.

When estimating variances we use the inverse of the estimated precision (17). While it may seem more natural to use the posterior mean of 
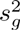
 as a point estimate for 
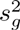
, we found that this can be very sensitive to small changes in the estimated hyperparameters **a**, and so can perform poorly. And while it may also be more natural to estimate variances on a log scale, for example using the posterior mean for log(*s_g_*), the absence of closed-form expressions makes this less convenient.

### 2.5 Unimodal prior assumption on variance or precision

The above formulation is based on assuming a unimodal prior distribution for the variance 
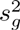
, and specifically by using a mixture of inverse-gamma distributions all with the same mode. An alternative is to assume a unimodal prior distribution for the precision 
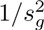
, by using a mixture of gamma distributions, all with the same mode. This is equivalent to using a mixture of inverse-gamma distributions for the variance 
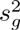
 as in (3) above, but with

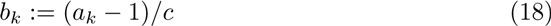

in place of (4), because the mode of a Gamma(*a*, *b*) distribution is at *c* = (*a* − 1)/*b*. We present results for both approaches. In practice one can assess which of the two models provides a better fit to the data by comparing their (maximized) likelihoods (12). Note that in many (but not all) cases the fitted prior distributions under either or both approaches will end up being unimodal for both the variance *and* the precision. However, even in these cases, the optimal likelihood under each approach will typically differ because the family of unimodal distributions being optimized over is different.

## 3 Results

### 3.1 Simulation studies

To compare and contrast our method with limma and limmaR we simulate data from the model (1)-(3), with *G* = 10,000, and degrees of freedom *df* = 3,10, 50 (corresponding to sample sizes 4, 11 and 51 respectively) under various scenarios for the actual distribution of variances (scenarios A-D) or precisions (scenarios E-H), as summarized in Tables 1 and 2. The simulation scenarios are designed to span the range from a single inverse-gamma prior as assumed by limma, to more complex distributions under which we might expect our method to outperform limma. Specifically we consider:

- Single IG (Gamma): single component inverse-gamma prior on variance (or gamma prior on precision), which satisfies the assumptions of limma.
- Single IG (Gamma) with outliers: two component inverse-gamma prior on variance (or gamma prior on precision), where one component models the majority of genes and the other component, being more spread out, attempts to capture possible outliers. The method limmaR is specifically designed to deal with the case where large variance outliers exist.
- IG (Gamma) mixture: a more flexible inverse-gamma mixture prior on variance (or mixture gamma prior on precision) with multiple components.
- Long tail log-normal mixture: log-normal mixture prior on variance or precision, which yields a longer tail than either the inverse-gamma or the gamma distribution.

**Table 1:**
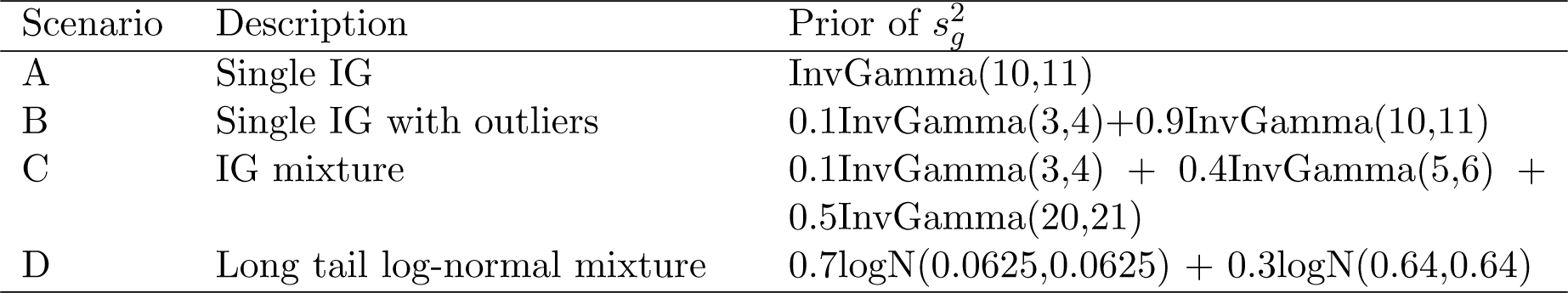
1Parameters for the simulation scenarios with unimodal prior on variance

**Table 2:** Parameters for the simulation scenarios with unimodal prior on precision

| Scenario | Description | Prior of $1/s_g^2$ |
| --- | --- | --- |
| E | Single gamma | Gamma(10,9) |
| F | Single gamma with outliers | 0.1Gamma(2,1)+0.9Gamma(10,9) |
| G | Gamma mixture | 0.1Gamma(2,1) + 0.4Gamma(5,4) + 0.5Gamma(30,29) |
| H | Long tail log-normal mixture | 0.7logN(0.0625,0.0625) + 0.3logN(0.64,0.64) |

For each simulation scenario we simulate 50 datasets and apply limma, limmaR, and our proposed method (vash) to estimate 
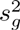
 (or 
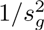
). We compare the relative root mean squared errors (RRMSEs) of the shrinkage estimators, which we define by

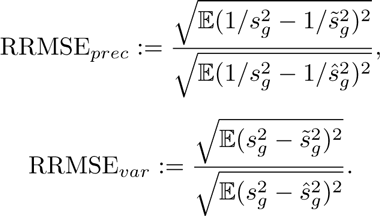

The RRMSE measures the improvement of a shrinkage estimator over simply using the sample variance 
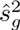
 or precision 
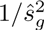
v, with RRMSE=1 indicating no benefit of shrinkage.

Figure 1 and 2 show the RRMSEs of limma, limmaR and vash for all scenarios. We summarize the main patterns as follows:

1. Across all scenarios, the mean RRMSE of vash is consistently no worse than either limma or limmaR, and is sometimes appreciably better. In contrast, limmaR sometimes performs better than limma and sometimes worse. In this sense vash is the most robust of the three methods.
2. In simulations under the simplest scenario (A and E) where the assumptions of limma are met, all three methods perform similarly. In particular, the additional flexibility of vash does not come at a cost of a drop of performance in the simpler scenarios.
3. When sample sizes are small (df=3) all methods perform similarly under all scenarios. This highlights the fact that the benefits of more flexible methods like vash are small if samples sizes are too small to exploit the additional flexibility. Put another way, for small sample sizes simple assumptions suffice.
4. When sample sizes are large (df=50) vash can outperform the other methods, particularly under the more complex scenarios (C,D; G,H), which most strongly violate the assumptions of limma. Indeed, in these cases both limma and limmaR can have RRMSE>1, indicating that they perform worse than the unshrunken sample estimators. That is, when sample sizes available to estimate each variance are relatively large shrinkage estimates based on oversimplified assumptions can make estimation accuracy worse rather than better. (In contrast, for small sample sizes, the benefits of shrinkage greatly outweigh any cost of oversimplified assumptions.)

**Figure 1:**
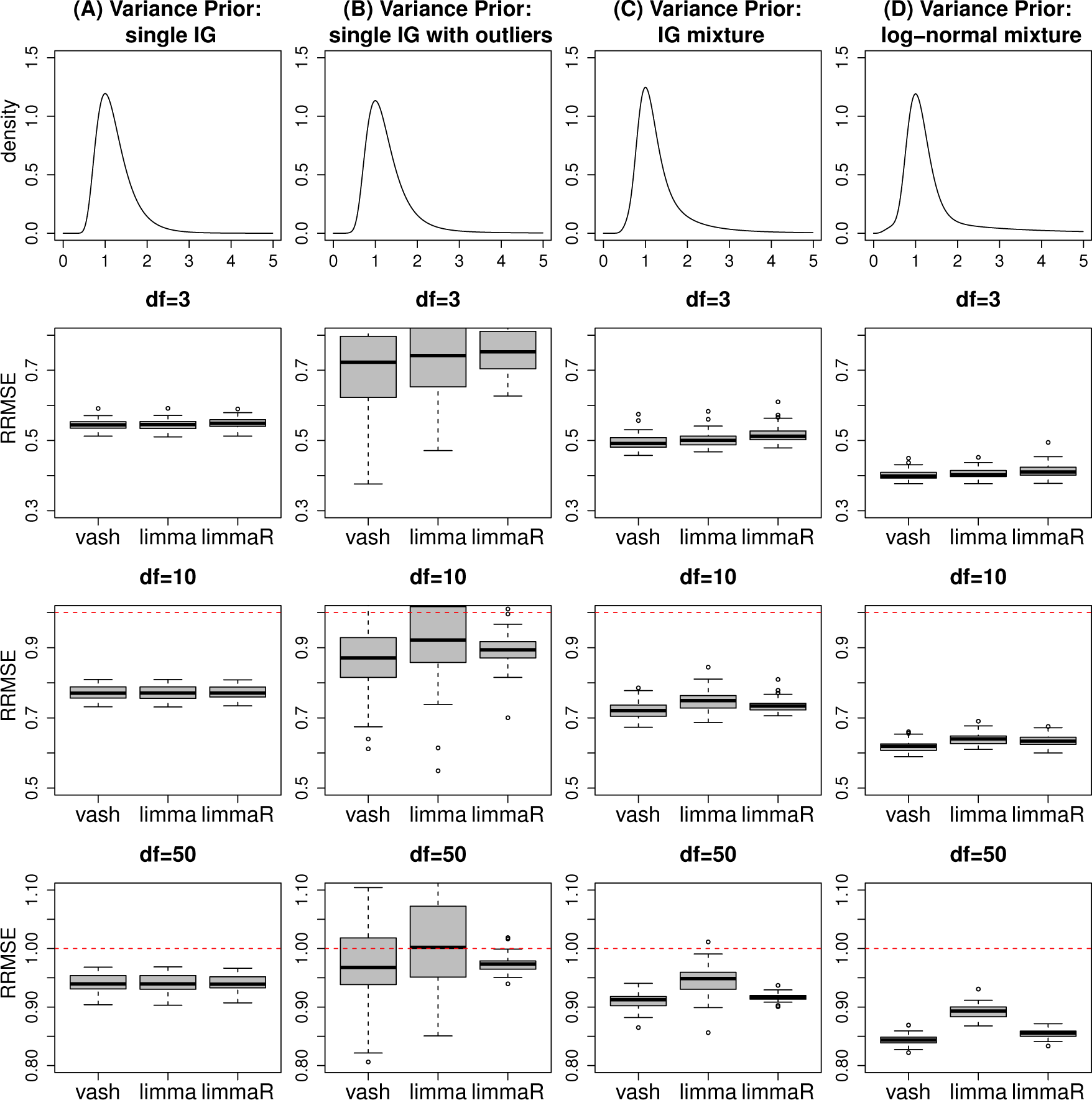
RRMSE*_var_* of three gene-specific variances estimators, limma, robust limma (limmaR) and our proposed estimator (vash) in the 4 simulation scenarios A-D with unimodal variance prior.

**Figure 2:**
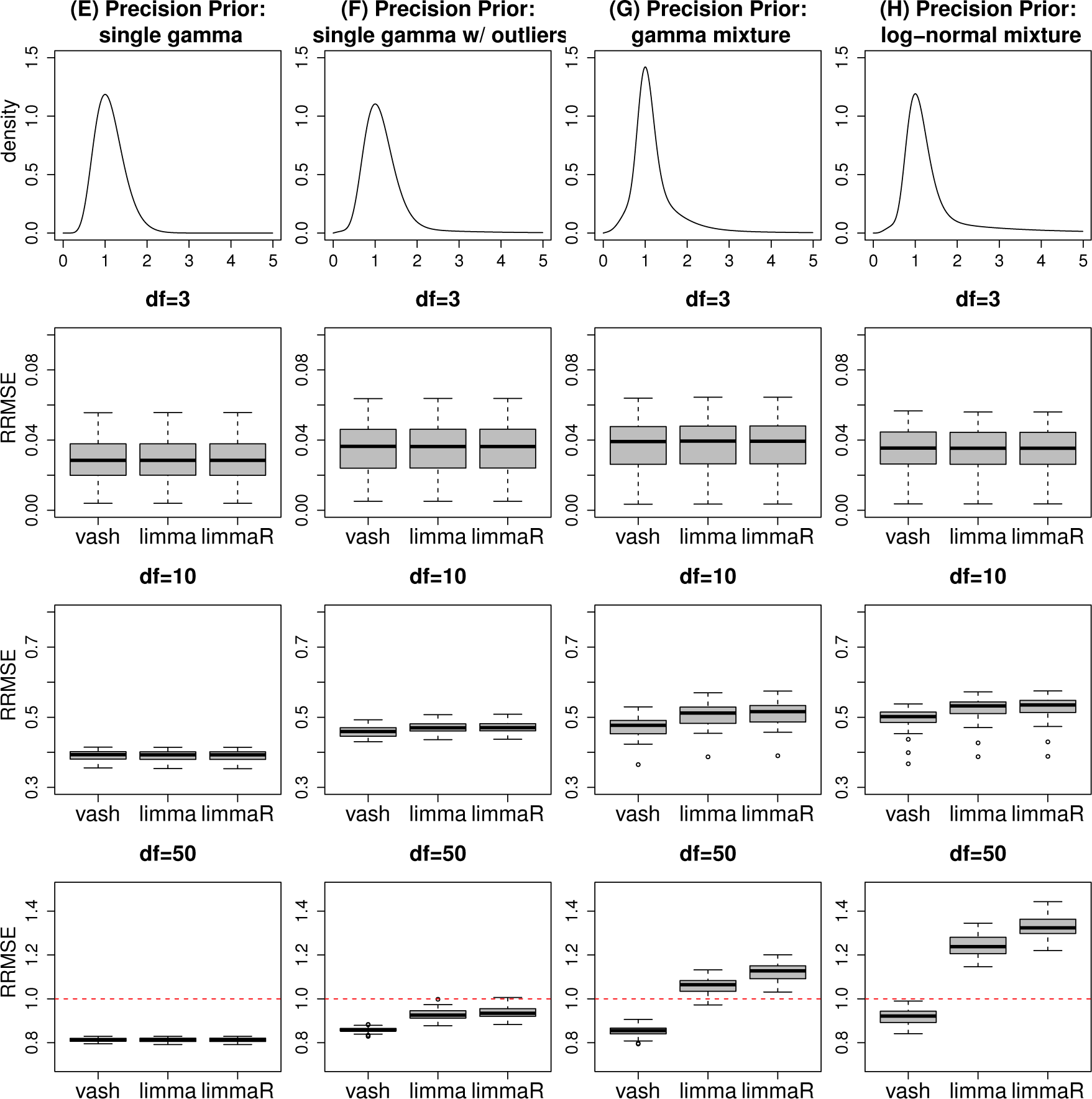
RRMSE*_prec_* of three gene-specific variances estimators, limma, robust limma (limmaR) and our proposed estimator (vash) in the 4 simulation scenarios E-H with unimodal precision prior.

We also note that in scenario B where variances are sampled from a two component inverse-gamma mixture prior (one “majority” component and one “outlier” component), both vash and limmaR perform similarly on average (and slightly outperform limma), but results of vash are slightly more variable than limmaR. Possibly this reflects the fact that limmaR was specifically designed to deal with such cases.

### 3.2 Assessment of variances in gene expression data

The results above demonstrate that the more flexible mixture prior implemented in vash, can in principle provide more accurate variance and precision estimates than the simple inverse-gamma prior implemented in limma. However, in practice these gains will only be realized if the actual distribution of variances differs from the single inverse-gamma model. Here we examine this issue using RNA sequencing data from the Genotype-Tissue Expression (GTEx) project [Lonsdale *et al.*, 2013]. The GTEx Project is an extensive resource which studies the relationship among genetic variation, gene expression, and other molecular phenotypes in multiple human tissues. Here we consider RNA-seq data (GTEx V6 dbGaP accession phs000424.v6.p1, release date: Oct 19, 2015, http://www.gtexportal.org/home/) on 53 human tissues from a total of 8555 samples (ranging from 6 to 430 samples per tissues).

Since in practice variance estimation is usually performed as part of a differential expression analysis [Smyth, 2004], we mimicked this set-up here: specifically we considered performing a differential expression analysis between every pair of tissues, estimating the variances *ŝ* in each case for the top 20,000 highly expressed genes. Since there are 53 tissues this resulted in 1378 datasets of variance estimates.

First, for each data set, we quantified the improved fit of the mixture prior vs a single component prior by comparing the maximum log-likelihood under each prior. (For the mixture prior we fitted both the unimodal-variance and unimodal-precision priors, and took the one that provided the larger likelihood.) In principle the mixture prior log-likelihood should always be larger because it includes the single component as a special case; we observed rare and minor deviations from this in practice due to numerical issues. Across all 903 datasets the average gain in log-likelihood of the mixture prior vs the single component prior was 34.1. The 25% quantile, median, 75% quantile, 90% quantile and maximum of the difference are given by 2.9, 15.8, 42.9, 77.4 and 705.2 respectively. A log-likelihood difference of 15.8 is already quite large: for comparison the maximum difference in log-likelihood for simulations under a single component model, Scenario A, df=50, was 1.9. We therefore conclude that the mixture component prior fits the data appreciably better for many datasets.

To visualize the deviations from a single component prior present in these data, we examine the fitted priors in datasets where the log-likelihood differences are about 42.9 (75% quantile), 77.4 (90% quantile) and higher. Figure 3 compares the fitted single component prior and mixture prior on several typical scenarios. Generally, the mixture priors use extra components to better fit the middle portion of distribution. The single component priors can match the tails pretty well, but often fails to accurately capture the peak.

**Figure 3:**
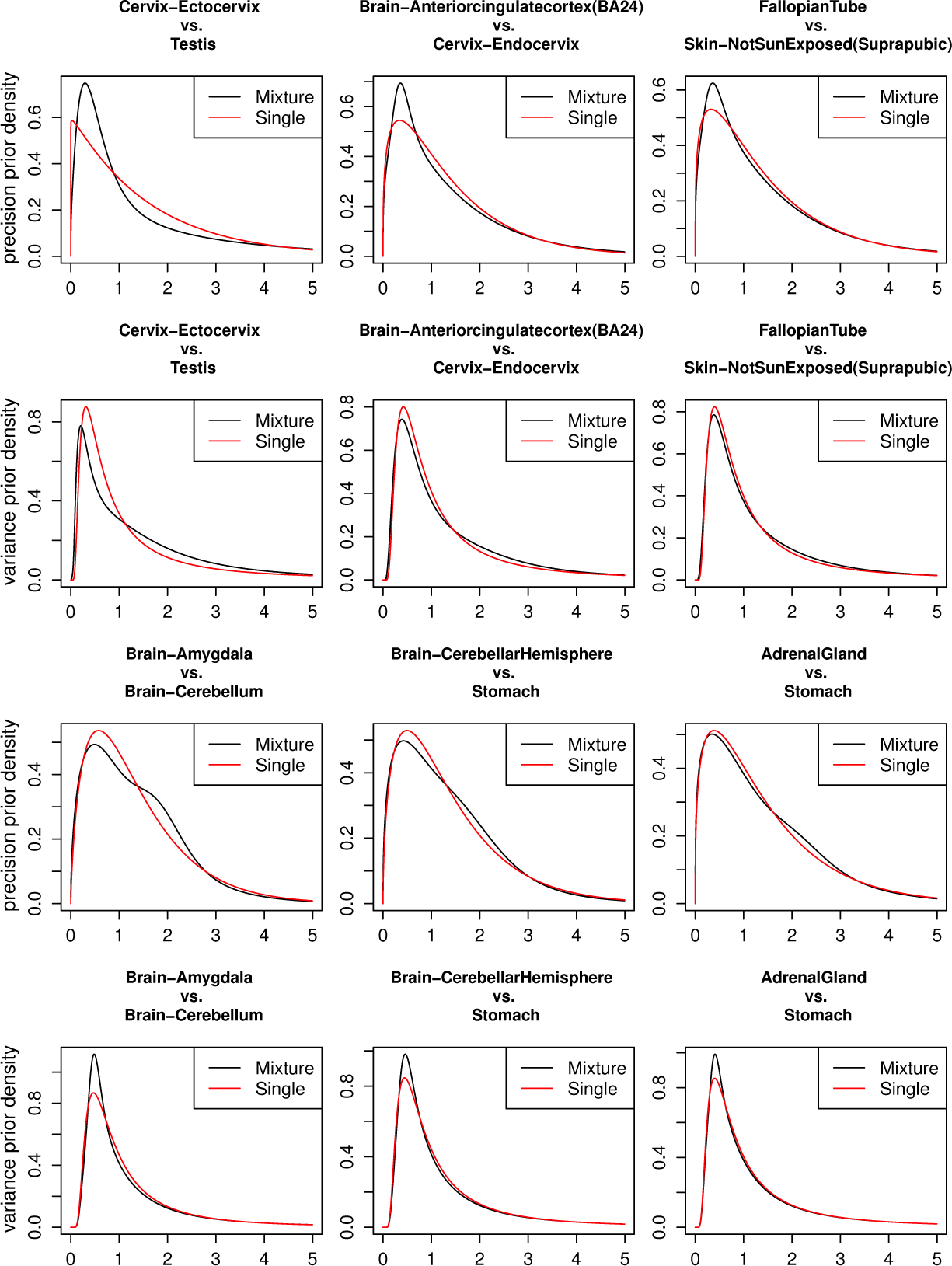
The variance priors (the 2nd and 4th row) and precision priors (the 1st and 3rd row) fitted by mixture prior model (black line) or single component prior model (red line) for 6 tissue pair comparisons. The differences in the log-likelihood between the mixture prior model and the single component prior model for tissue pair comparisons “Cervix-Ectocervix vs Testis”, “Brain-Amygdala vs Brain-Cerebellum”, “Brain-Anteriorcingulatecortex (BA24) vs Cervix-Endocervix”, “Brain-CerebellarHemisphere vs Stomach”, “Fallopian Tube vs Skin-Not Sun Exposed (Suprapubic)”, “Adrenal Gland vs Stomach” are given by 705, 166, 78, 78, 44, 44 respectively (from top-left to bottom-right).

Overall, our impression from Figure 3 is that differences between the fitted priors seem relatively minor, and might be expected to lead to relatively small differences in accuracy of shrinkage estimates, despite the large likelihood differences. To check this impression we simulated data where the variances are generated from the fitted mixture priors for four of these datasets (the four datasets on the right hand side of Figure 3). Figure 4 compares the RRMSEs of vash, limma and limmaR in these four scenarios. In general the results confirm our impression: the three methods perform very similarly in most scenarios, although vash shows some gain in accuracy in two scenarios with df=50.

**Figure 4:**
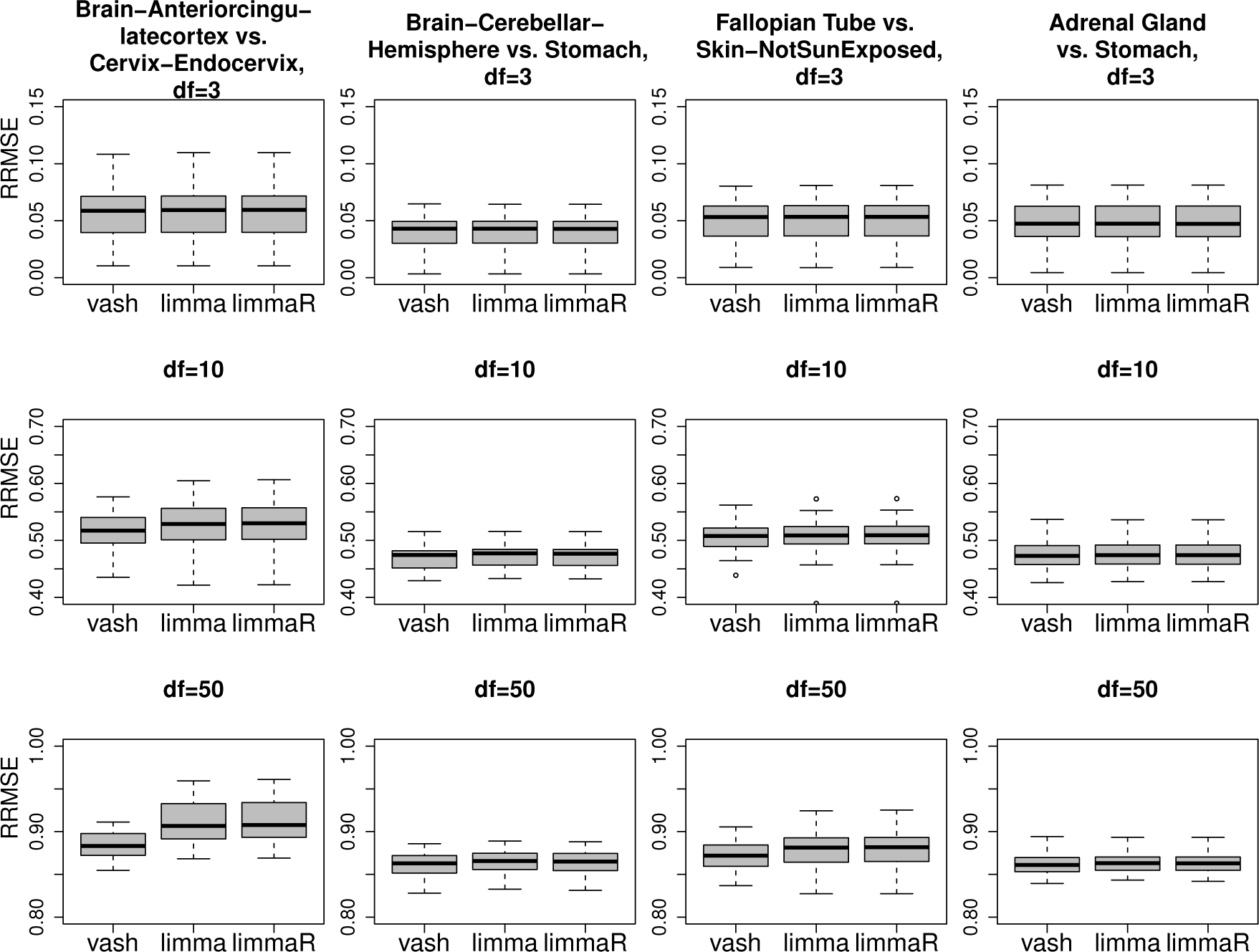
RRMSE_prec_ of three gene-specific variances estimators, limma, robust limma (limmaR) and our proposed estimator (vash) in simulation scenarios, which simulate the last four GTEx tissue pair comparisons (“Brain-Anteriorcingulatecortex (BA24) vs Cervix-Endocervix”, “Brain-CerebellarHemisphere vs Stomach”, “Fallopian Tube vs Skin-Not Sun Exposed (Suprapubic)” and “Adrenal Gland vs Stomach”) in Figure 3.

## 4 Discussion

We have presented a flexible empirical Bayes approach (“variance adaptive shrinkage”, or “vash”) to shrinkage estimation of variances. The method makes use of a mixture model to allow for a flexible family of unimodal prior distributions for either the variances or precisions, and uses an accelerated EM-based algorithm to efficiently estimate the underlying prior by maximum likelihood. Although slower than limma, vash is computationally tractable for large datasets: for example, for data with 10,000 genes, vash typically takes about 30 seconds (limma takes just a few seconds).

Our results demonstrate that vash provides a robust and effective approach to variance shrinkage, at least in settings where the distribution of the variances (or precisions) is unimodal. When the true variances come from a single inverse-gamma prior, vash is no less accurate than the simpler method. When the variances come from a more complex distribution vash can be more accurate than simpler methods if the sample sizes to estimate each variance are sufficiently large.

In the gene expression datasets we examined here, the gains in accuracy of vash vs limma are small, and likely not practically important. While this could be viewed as disappointing, it nonetheless seems useful to show this, since it suggests that in many gene expression contexts the simpler approaches will suffice. At the same time, it remains possible that our method could provide practically useful gains in accuracy for other data-sets, and as we have shown, it comes at little cost. In addition, our work provides an example of a general approach to empirical Bayes shrinkage - use of mixture components with a common mode to model unimodal prior distributions that could be useful more generally.

Our method is implemented in an R package **vashr** available from http://github.com/mengyin/vashr. Codes for reproducing analyses and figures in this paper are athttps://github.com/mengyin/vash.

## Acknowledgements

We thank the the NIH GTEx project for providing RNA-seq datasets.

## Appendix

Details of the algorithm used to estimate hyper-parameters *c* and *π*:

To maximize the likelihood (12) we iteratively update *c* and *π* using the following steps until they converge:

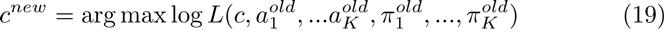

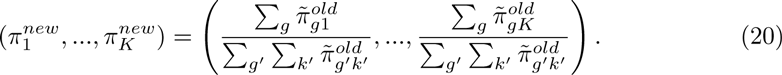

Because there is no analytic solution to the optimization problem (19) we approximate the optimum via the quasi-Newton method using the function optim in R. The updating procedure (19)-(20) is a fixed point mapping function, hence we can accelerate the overall optimization procedure by the R package SQUAREM [Varadhan, 2010], which is computationally efficient in solving fixed point problems.

